# The reward-complexity trade-off in schizophrenia

**DOI:** 10.1101/2020.11.16.385013

**Authors:** Samuel J. Gershman, Lucy Lai

## Abstract

Action selection requires a policy that maps states of the world to a distribution over actions. The amount of memory needed to specify the policy (the policy complexity) increases with the state-dependence of the policy. If there is a capacity limit for policy complexity, then there will also be a trade-off between reward and complexity, since some reward will need to be sacrificed in order to satisfy the capacity constraint. This paper empirically characterizes the trade-off between reward and complexity for both schizophrenia patients and healthy controls. Schizophrenia patients adopt lower complexity policies on average, and these policies are more strongly biased away from the optimal reward-complexity trade-off curve compared to healthy controls. However, healthy controls are also biased away from the optimal trade-off curve, and both groups appear to lie on the same empirical trade-off curve. We explain these findings using a cost-sensitive actor-critic model. Our empirical and theoretical results shed new light on cognitive effort abnormalities in schizophrenia.

## Introduction

People diagnosed with schizophrenia are typically less willing to exert cognitive and physical effort to obtain rewards (Culbreth et al., 2018). For example, Culbreth et al. (2016) gave subjects the opportunity to earn more reward by exerting greater effort (choosing higher working memory loads in the N-back Task). Compared to healthy controls, schizophrenia patients exhibited a greater preference for low effort / low reward tasks, and the strength of this preference correlated with negative symptom severity. Similar results have been reported using other assays of cognitive effort (Fortgang et al., 2020, Reddy et al., 2015, Wolf et al., 2014), although the literature is inconsistent (Gold et al., 2015, Horan et al., 2015).

One obstacle to a unified understanding of cognitive effort abnormalities in schizophrenia is the heterogeneity of the constructs.^1^ For example, the Deck Choice Effort Task used in Horan et al. (2015) operationalizes cognitive effort in terms of task switching (greater effort for more frequent switches). The Demand Selection Task (Kool et al., 2010) used by Gold et al. (2015) similarly manipulates cognitive effort by varying task switching frequency. Both studies failed to find changes in cognitive effort avoidance related to schizophrenia. A large-scale transdiagnostic assessment using the Demand Selection Task also found no relationship between sub-clinical schizotypy and cognitive effort avoidance (Patzelt et al., 2019). These results suggest that the representation of *computational cost* may be unaffected in schizophrenia.

The N-back task, in contrast, is effortful in the sense that it taxes representational resources needed for storing information in memory. In other words, it incurs an *informational cost* that can be formalized using information theory (Brady et al., 2009, Miller, 1956, Sims et al., 2012, Sims, 2016). We can think of the memory system as a communication channel that encodes a stream of stimuli into codewords, and then decodes the stimuli from these codewords at the time of retrieval. If the encoder is noisy, then the decoder will make errors. To reduce this error, the encoder can use longer codewords that store information redundantly (analogous to how you might repeat something multiple times to make sure another person heard you). If the encoder is capacity-limited (the code length cannot exceed some bound), then there is a limit to how much it can reduce its error (Shannon, 1948). This information-theoretic framework gives us a precise way of talking about the nature of cognitive effort in working memory tasks: increasing the code length for information storage is effortful. The evidence from the incentivized N-back Task (Culbreth et al., 2016) suggests that schizophrenia patients are less willing to pay informational effort costs. We pursue this hypothesis further using a different task and a theoretical framework that makes the informational costs explicit. Collins and Frank (2012) introduced a reinforcement learning task in which subjects selected one of 3 actions on each trial and received reward feedback that depended on a trial-specific stimulus (Figure 1). The number of distinct stimuli (the set size) was manipulated across blocks. Performance decreased as a function of set size, which the authors interpreted in terms of a capacity-limited working memory contribution to reinforcement learning. Using this task, Collins and colleagues (Collins et al., 2017a, 2014) found that schizophrenia patients also exhibited a set size effect, but with overall lower performance, consistent with the hypothesis that the patients had lower working memory capacity for reinforcement learning. This finding agrees with an established literature on working memory impairments in schizophrenia (Lee and Park, 2005).

**Figure 1:**
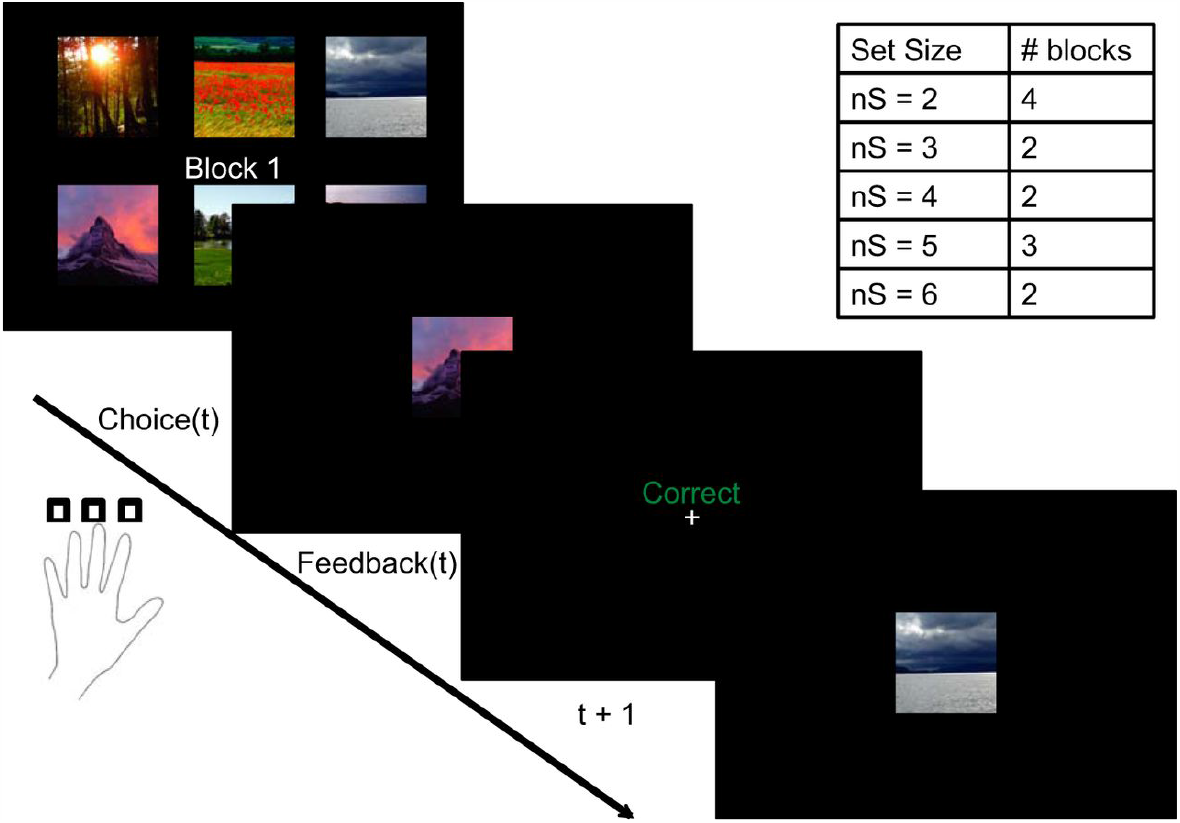
Task schematic. On each trial, subjects selected one of three different actions conditional on a presented stimulus, and were then presented with deterministic reward feedback (correct/incorrect). The number of stimuli (the set size) varied across blocks.

Gershman (2020) analyzed data from the Collins task through the lens of rate distortion theory (Berger, 1971), which addresses the interface between information theory and statistical decision theory. Following earlier work (Parush et al., 2011, Still and Precup, 2012, Tishby and Polani, 2011), informational costs were defined in terms of *policy complexity*—the mutual information between states and actions (explained further below). Intuitively, policy complexity measures the amount of memory required to specify a policy mapping states to actions. If the policy is highly state-dependent (e.g., a look-up table), then the memory required will be high, compared to a policy that is relatively state-independent (actions do not depend on states). If there is a bound on policy complexity, then there will be a trade-off between reward and complexity: some reward must be sacrificed in order to satisfy the complexity bound. This gives rise to a form of perseveration, the tendency to produce the same action policy across states regardless of the reward outcome. In this paper, we apply the same analyses used in Gershman (2020) to data from schizophrenia patients, in order to characterize their reward-complexity trade-off.

A key goal of this paper is to understand to what extent differences in cognitive effort between patients and controls, as well as differences between individuals within these groups, can be understood as a rational trade-off. Specifically, an individual may choose to avoid cognitive effort based on their subjective preference for reward relative to the effort cost. Observing that schizophrenia patients exert less effort does not allow us to say whether they perceive cognitive effort as more costly relative to reward, or whether they are failing to optimize the trade-off between reward and effort. In the latter case, schizophrenia patients may in fact be willing to exert more effort, but they fail to identify their subjectively optimal level of effort. Rate distortion theory provides us with the theoretical tools to address how close schizophrenia patients and healthy controls are to the optimal reward-complexity trade-off. If they adhere closely to the optimal trade-off curve, then we have a basis for claiming that any differences in policy complexity between the two groups reflects a rational trade-off.

## Methods and Materials

### Theoretical framework

We model an agent that visits states (denoted by *s*) and takes actions (denoted by *a*). We assume that the agent learns a *value function Q*(*s, a*) that defines the expected reward in state *s* after taking action *a*. In the experiment analyzed here, the value function is deterministic, so in principle it can be learned in a few trials, or even a single trial. For simplicity, we treat the value function as known; even though this is not an accurate characterization of the learning process (see next section), we expect that it will adequately capture the average behavior of subjects, which is our focus here. Each state is visited with probability *P* (*s*), and an action is chosen according to a policy *π*(*a* | *s*). The average number of bits (or *rate*) necessary to encode a policy with arbitrarily small error is equal to the mutual information between states and actions:

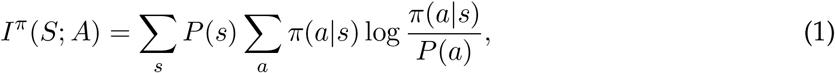

where *P* (*a*) = Σ_*s*_ *P* (*s*)*π*(*a*| *s*) is the marginal probability of choosing action *a* (i.e., the policy averaged across states). Because the mutual information quantifies the degree of probabilistic dependency between states and actions, we will refer to it as the *policy complexity*. State-dependent policies are more complex than state-independent policies. Thus, policy complexity is minimized (mutual information is equal to 0) when the policy is the same in every state.

The agent’s goal is to earn as much reward as possible, subject to the constraint that the policy complexity cannot exceed a capacity limit. Formally, the resource-constrained optimization problem is defined as follows:

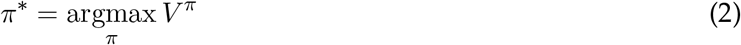

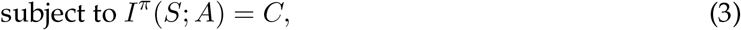

Where

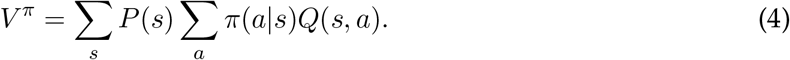

is the average reward under policy *π*, and *C* is the channel capacity—the maximum achievable policy complexity. Two other necessary constraints (action probabilities must be non-negative and sum to 1) are left implicit. This constrained optimization problem can be equivalently expressed in a Lagrangian form:

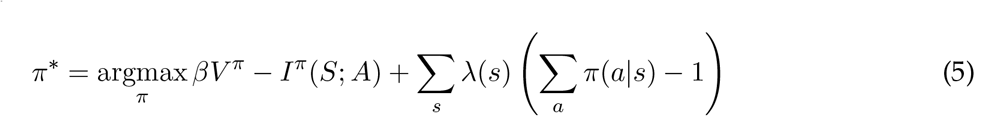

with Lagrange multipliers *β* and *λ*(*s*). The optimal policy *π*^*∗*^ has the following form (Parush et al., 2011, Still and Precup, 2012, Tishby and Polani, 2011):

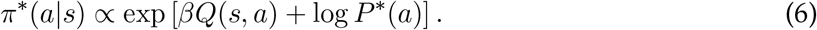

The optimal policy thus takes the form of a softmax function, with a frequency-dependent bias (perseveration) term. The Lagrange multiplier *β* plays the role of the “inverse temperature” parameter, which regulates the exploration-exploitation trade-off via the amount of stochasticity in the policy (Sutton and Barto, 2018). When *β* is close to 0, the policy will be near-uniform, and as *β* increases, the policy will become increasingly concentrated on the action with maximum value. The inverse of *β* is the partial derivative of the value with respect to the policy complexity:

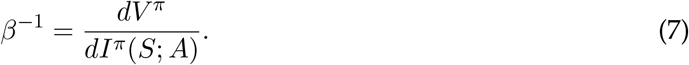

Geometrically, this is the slope of the optimal reward-complexity curve for a particular resource constraint (see below).

The perseveration term implicitly depends on the optimal policy:

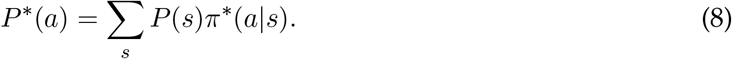

To find the optimal policy, we can use a variation of the *Blahut-Arimoto algorithm* (Arimoto, 1972, Blahut, 1972), alternating between updating the policy according to Eq. 6 and updating the marginal action distribution according to Eq. 8. By performing this optimization for a range of *β* values, we can construct a reward-complexity curve that characterizes the optimal policy for a given resource constraint.

### A process model: cost-sensitive actor-critic learning

The previous section presented a computational-level account of policy optimization under an information-theoretic capacity limit. For convenience, we assumed direct access to the reward function, and computed the optimal policy using the Blahut-Arimoto algorithm. However, these idealizations are not plausible as process models. Real agents need to learn the reward function from experience, and the Blahut-Arimoto algorithm may be computationally intractable when the state space is large (because it requires marginalization over all states according to Eq. 8).

To derive a more cognitively plausible process model, we start from the observation that the Lagrangian optimization problem in Eq. 5 can be expressed in terms of an expectation over states:

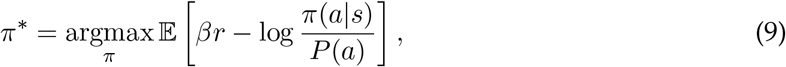

This formulation allows us to construct an “actor-critic” learning rule using the stochastic policy gradient algorithm (Sutton and Barto, 2018), which directly optimizes Eq. 9 by taking the gradient of the average reward with respect to the policy parameters. First, we define a parametrized policy (the “actor”):

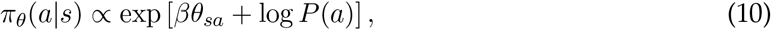

where *θ* denotes the policy parameters. Note that this parametrization mirrors the optimal parametrization in Eq. 6. Given an observed reward *r* after taking action *a* in state *s*, the policy parameters are updated according to:

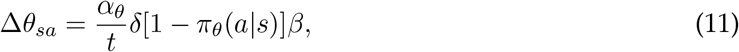

where *α*_*θ*_ is the actor learning rate, *N* is the set size, and

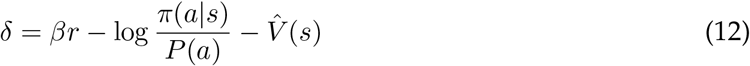

is the prediction error of the “critic” 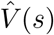, an estimator of the expected cost-sensitive reward, updated according to:

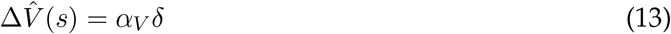

with critic learning rate *α*_*V*_. We scaled the actor learning rate (but not the critic learning rate) by 1*/t* for two reasons. This ensures that the the policy eventually converges to the optimal policy by satisfying the Robbins-Munro conditions for stochastic approximation algorithms (Robbins and Monro, 1951), and by ensuring that the actor learning rate will generally be slower than the critic learning rate (Konda and Tsitsiklis, 2000).

To complete the model, we estimate the marginal action probabilities with an exponential moving average:

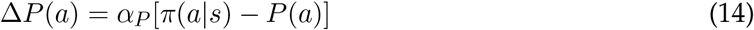

with learning rate *α*_*P*_.

We fit four free parameters (*β, α*_*θ*_, *α*_*V*_, *α*_*P*_) to each individual’s choice behavior using maximum likelihood estimation. To assess the match to the data, we then simulated the fitted model for each participant, using the same stimuli presented to the human subjects.

### Data set

We applied the theory to a data set originally reported in Collins et al. (2014). Subjects performed a reinforcement learning task in which the set size (the number of distinct stimuli, corresponding to states) varied across blocks (Figure 1). On each trial, subjects saw a single stimulus, chose an action and received deterministic reward feedback. Each stimulus was associated with a single rewarded action. Each subject completed 13 blocks, with set sizes ranging from 2 to 6. Each stimulus appeared 9-15 times in a block, based on a performance criterion of at least 4 correct responses of the last 5 presentations of each stimulus. No stimulus was repeated across blocks.

Two groups of subjects (schizophrenia patients and healthy controls) completed the experiment. The schizophrenia group (henceforth denoted SZ) consisted of 49 people (35 males and 14 females) with a DSM-IV diagnosis of schizophrenia (*N* = 44) or schizoaffective disorder (*N* = 5). The healthy control group (henceforth denoted HC) consisted of 36 people (25 males and 11 females), matched to the patient group in terms of demographic variables, including age, gender, race/ethnicity, and parental education.

### Estimating empirical reward-complexity curves

To construct the empirical reward-complexity curve, we computed for each subject the average reward and the mutual information between states and actions. From the collection of points in this two-dimensional space, we could estimate an empirical reward-complexity curve. While there are many ways to do this, we found 2nd-order polynomial regression to yield a good fit. To estimate mutual information, used the technique introduced by Hutter (2002), which computes the posterior expected value of the mutual information under a Dirichlet prior. Following Gershman (2020), we chose a symmetric Dirichlet prior with a concentration parameter *α* = 0.1, which exhibits reasonably good performance when the joint distribution is sparse (Archer et al., 2013). The sparsity assumption is likely to hold true in the data set analyzed here because there is a single rewarded action in each state.

## Results

### Comparing optimal and empirical reward-complexity curves

How close are subjects to the optimal reward-complexity trade-off curve? Figure 2A-E compares the optimal and empirical curves, broken down by set size and subject group. We can glean several insights from these plots. First, despite a gap between the optimal and empirical trade-off curves (explored further below), there was a strong correlation between the curves for both groups (*r* = 0.94 for HC, *r* = 0.92 for SZ, both *p <* 0.00001).^2^. This finding affirms earlier work (Gershman, 2020) showing that people approach the optimal reward-complexity trade-off, particularly for those exhibiting high policy complexity. Second, recapitulating findings from earlier work using variants of this task (Collins, 2018, Collins et al., 2017b, Collins and Frank, 2012, 2018), subjects earn less reward with larger set sizes, indicating a resource constraint on reinforcement learning. Third, average policy complexity did not vary monotonically across set sizes for either group (Fig. 2F), indicating a roughly constant resource constraint. This finding is consistent with the hypothesis that set size effects reflect reallocation of a fixed resource across multiple items (Ma et al., 2014). Fourth, policy complexity was significantly lower for the SZ group [mixed-effects ANOVA: *F* (1, 415) = 11.51, *p <* 0.001], and did not interact with set size (*p* = 0.14), indicating that the subjects in the SZ group were tapping fewer cognitive resources in this task.

**Figure 2:**
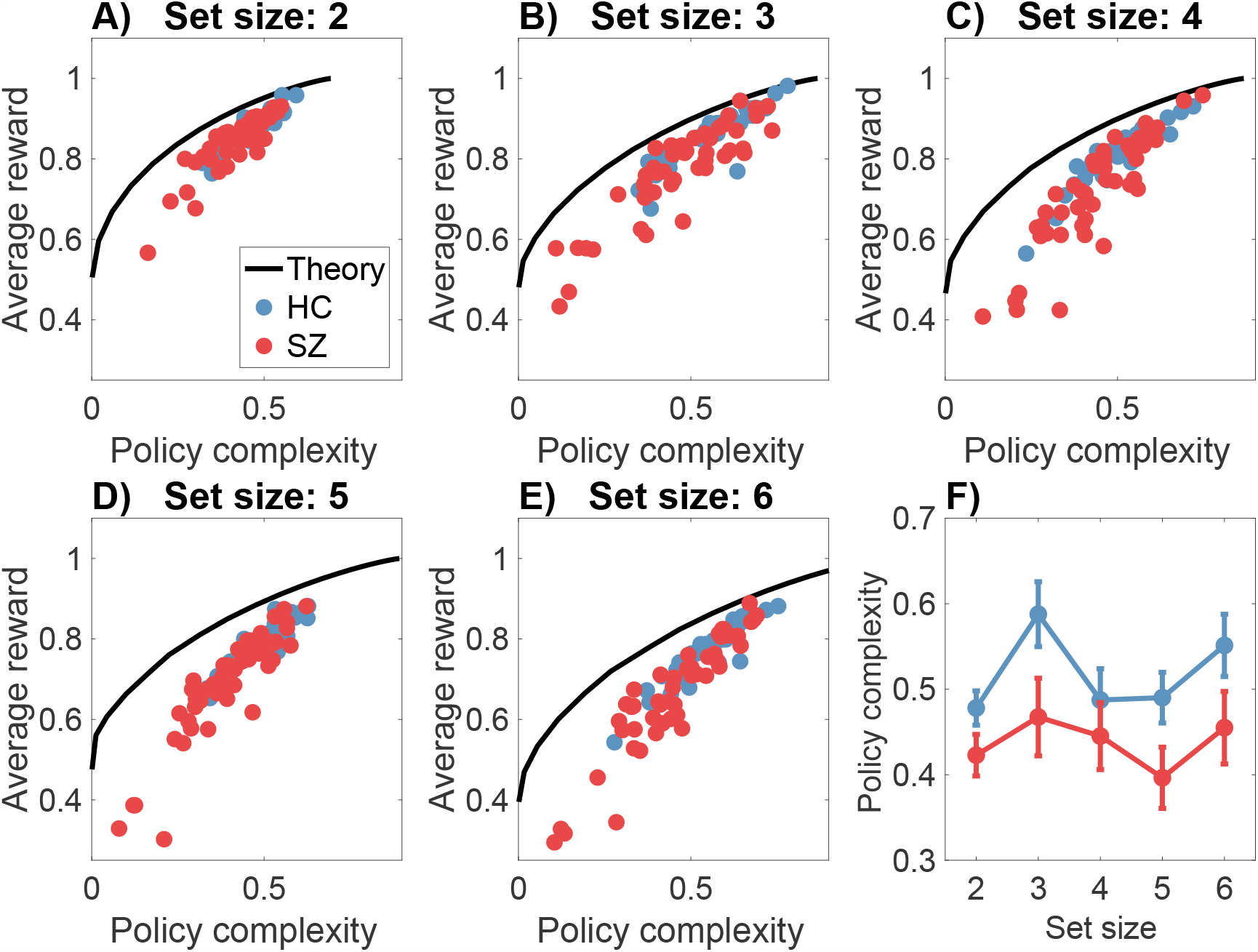
The reward-complexity trade-off. (A-E) Each panel shows the optimal reward-complexity curve (solid line) for a given set size, along with the empirical reward-complexity values (circles) for each subject (HC = healthy controls; SZ = schizophrenia patients). Policy complexity is measured in natural units of information (nats). A uniform distribution over actions corresponds to a policy complexity of 0. (F) Policy complexity as a function of set size. Error bars show 95% confidence intervals.

Figure 2 displays a systematic discrepancy between the optimal and empirical trade-off functions, which we quantify in terms of the *bias* (the difference between the two functions sampled at the empirical trade-off points). The average bias, broken down by set size and group, is shown in Figure 3A. A mixed-effects ANOVA found main effects of set size [*F* (4, 415) = 6.99, *p <* 0.001] and group [*F* (1, 415) = 5.76, *p <* 0.05], as well as an interaction [*F* (4, 415) = 4.07, *p <* 0.005]. Average bias was larger for higher set sizes and for the SZ group; the difference between the groups grew as a function of set size. Thus, subjects appear to deviate from optimality to a greater degree when cognitive demands are larger, and this deviation is exacerbated for SZ patients.

**Figure 3:**
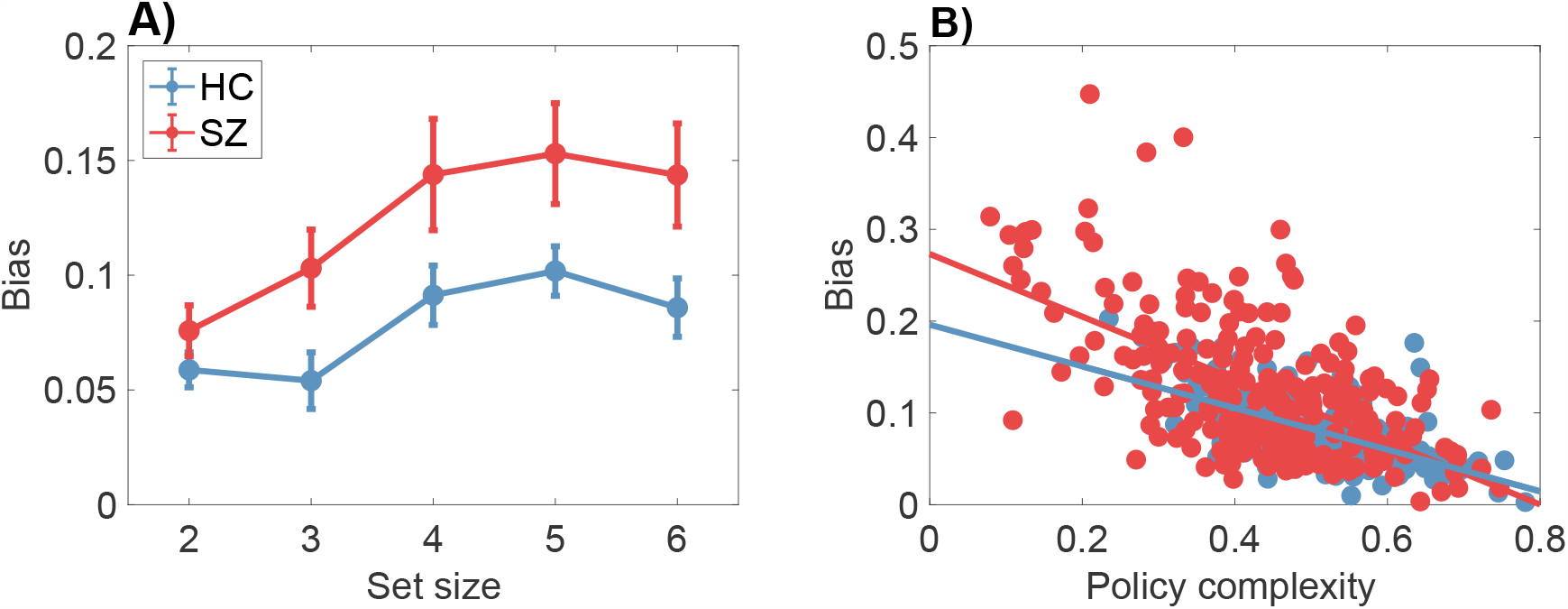
Bias differs between healthy control and schizophrenia patients. Bias is defined as the gap between the optimal and empirical reward-complexity curves. (A) Bias is larger for the schizophrenia group than for the healthy control group. Error bars show 95% confidence intervals. (B) Bias is negatively correlated with policy complexity. The correlation coefficient does not differ significantly between subject groups.

Another aspect of bias, captured in Figure 3B, is that it declines with policy complexity for both groups (Pearson correlation: *r* = *-* 0.62 for HC, *r* = *-* 0.61 for SZ, both *p <* 0.0001; Spearman correlation: *ρ* = *-* 0.60 for HC, *ρ* = *-* 0.57 for SZ, both *p <* 0.0001). In other words, subjects who have more cognitive resources available are closer to the optimal trade-off curve. Importantly, the Pearson correlation between bias and policy complexity did not differ significantly between the two groups (95% confidence interval for the correlation coefficient was [*-* 0.70, *-* 0.52] for HC and [*-* 0.68, *-* 0.53] for SZ). This indicates that the two groups, while differing in average bias, do not differ in their bias *functions*, an observation that dovetails with the analysis of empirical trade-off curves reported next.

We now turn to the critical question raised in the Introduction: do subjects in the two groups occupy different points along the same trade-off curve, or do they occupy different trade-off curves? To answer this question, we fit a parametric model (2nd-order polynomial regression) to the reward-complexity values, separately for the two groups and for each set size. This modeling demonstrated that the two groups have essentially the same trade-off curves. We show this in two ways. First, none of the parameter estimates differ significantly between groups for any of the set sizes (Figure 4A-E). Second, we compared the “independent” polynomial regression model, in which parameters are allowed to vary between the groups, to a “joint” model in which the parameters are forced to be the same (but still allowed to vary across set sizes). We compared models using the Bayesian information criterion (BIC), which applies a complexity penalty to the additional free parameters in the independent model. Across set sizes, the model comparison consistently favored the joint model (Figure 4F).

**Figure 4:**
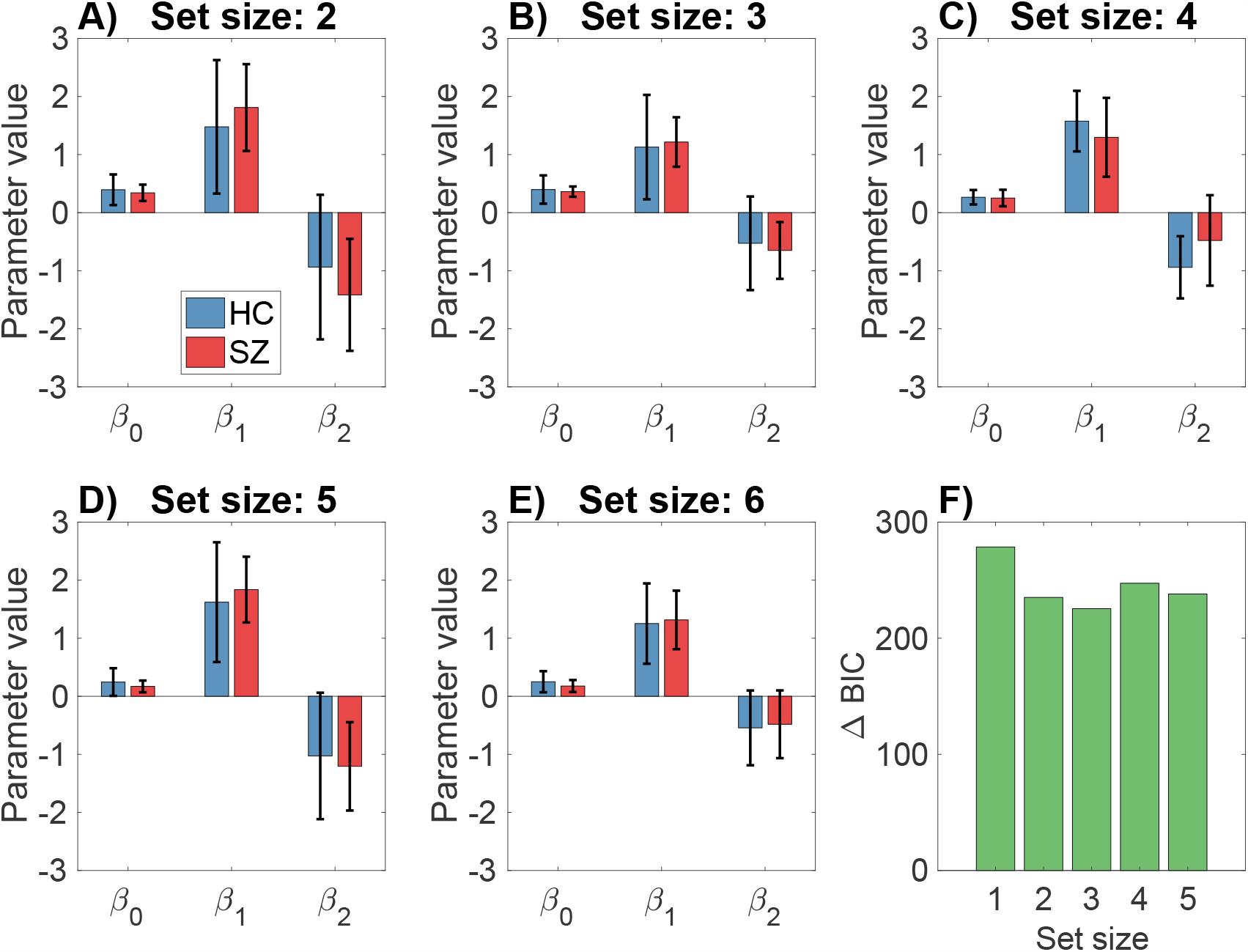
Polynomial regression modeling of empirical reward-complexity curves. (A-E) Regression coefficients do not differ between the healthy control and schizophrenia groups. Error bars show 95% confidence intervals. (F) The difference in the Bayesian information criterion (BIC) between the independent and joint regression models. Positive values favor the joint regression model.

### Modeling

As a first step towards understanding why the empirical and optimal trade-off curves diverge, we simulated a process model of policy optimization (see Materials and Methods). This model is a cost-sensitive version of the actor-critic model that has been studied extensively in neuroscience and computer science. The key idea is that the agent is penalized for policies that deviate from the marginal distribution over actions (i.e., the probability of taking a particular action, averaging over states). This favors less complex policies, because the penalty will be higher to the extent that the agent’s policy varies across states. Mechanistically, the model works like a typical actor-critic model, with the difference that the policy complexity penalty is subtracted from the reward signal. We fit the actor-critic model to the choice data using maximum likelihood estimation, and then simulated the fitted model on the task. Applying the same analyses to these simulations (Figures 5 and 6) verified that this model achieved a reasonably good match with the experimental data (compare to Figures 2 and 3), with the exception that it didn’t capture the empirically observed increase of bias with set size. We then asked to what extent different parameters contributed to the bias effect (i.e., the deviation between empirical and optimal trade-off curves). Entering the parameters for each subject into a linear regression with average bias as the dependent variable, we found significant positive coefficients for the actor learning rate (*α*_*θ*_, *t* = 2.49, *p <* 0.05; Figure 7B) and the marginal action probability learning rate (*α*_*P*_, *t* = 2.47, *p <* 0.05; Figure 7C).

**Figure 5:**
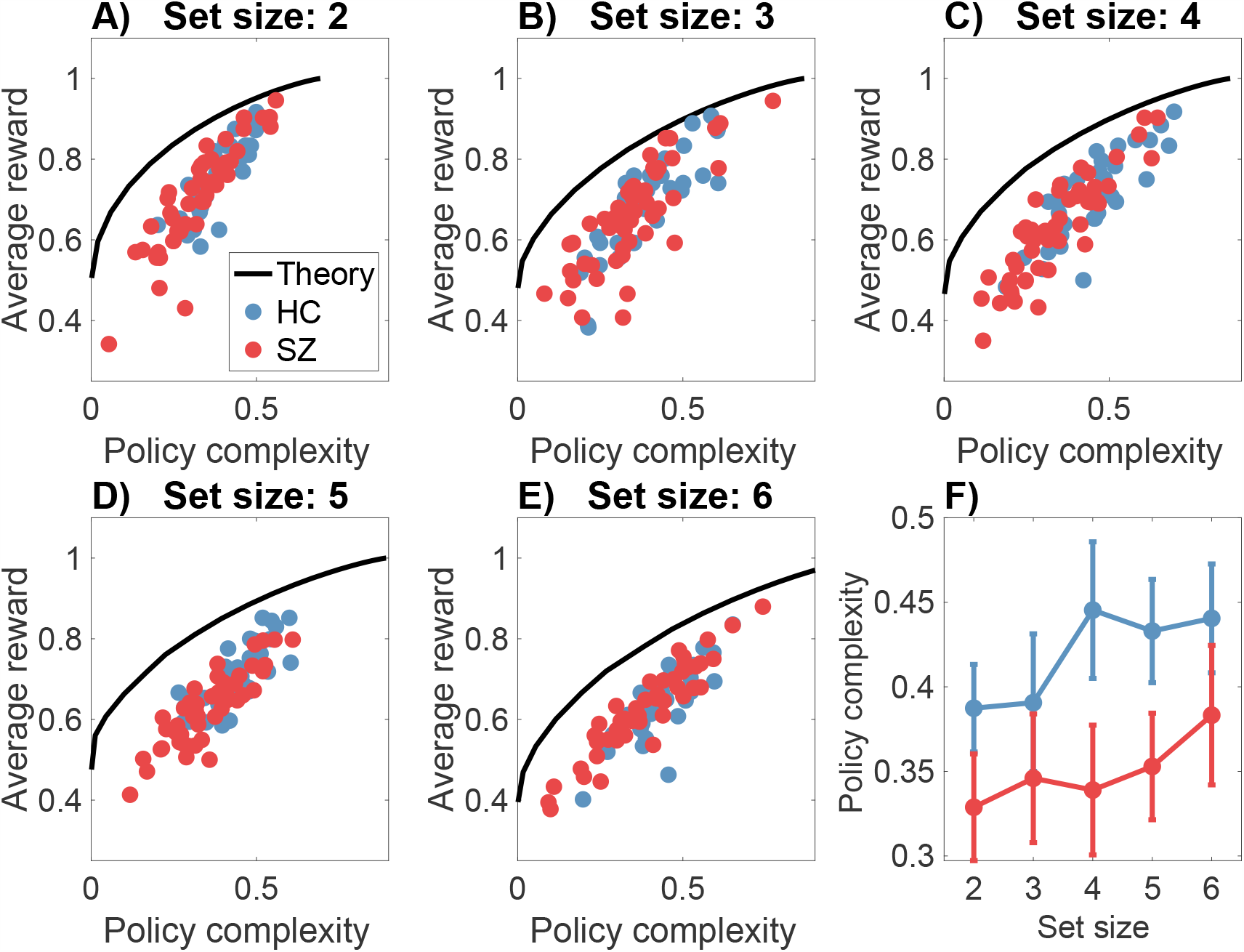
The reward-complexity trade-off for simulated cost-sensitive agents. (A-E) Each panel shows the optimal reward-complexity curve (solid line) for a given set size, along with the simulated reward-complexity values (circles) for each subject (HC = healthy controls; SZ = schizophrenia patients). (F) Policy complexity as a function of set size. Error bars show 95% confidence intervals.

**Figure 6:**
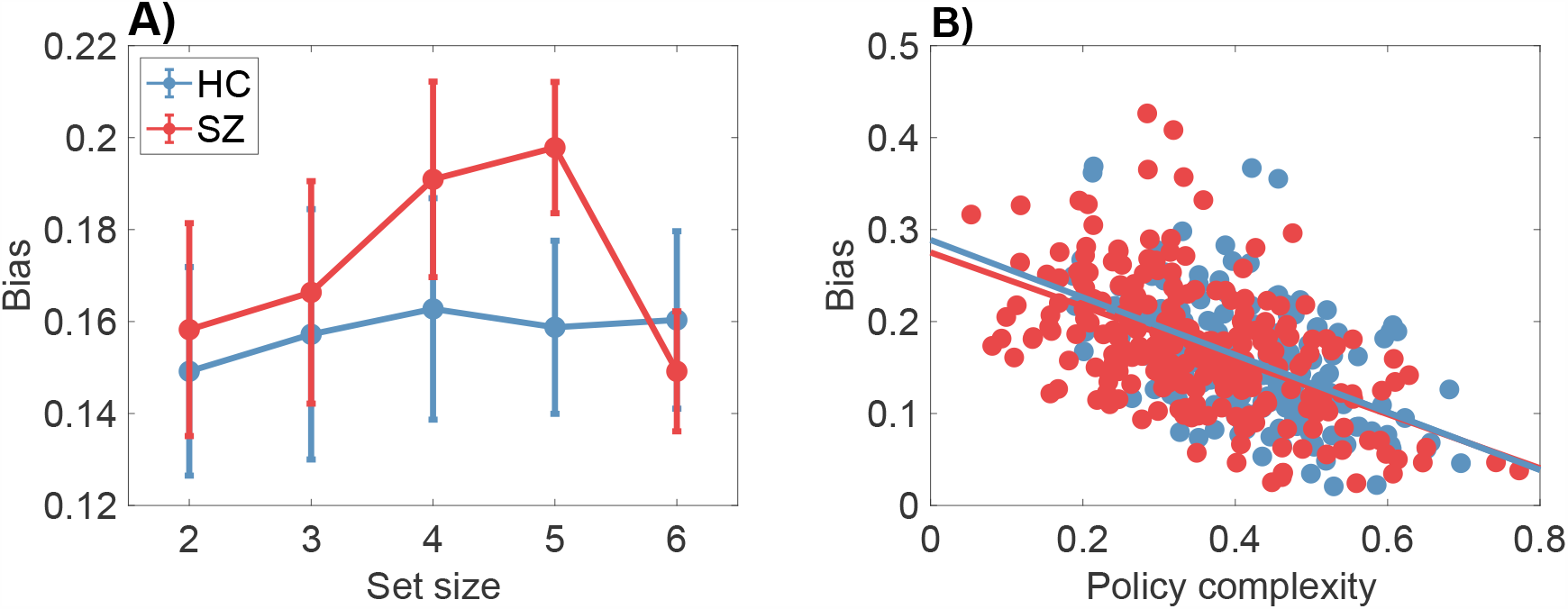
Bias differs between simulated healthy control and schizophrenia patients. (A) Bias is larger for the simulated schizophrenia group than for the healthy control group. Error bars show 95% confidence intervals. (B) Bias is negatively correlated with policy complexity. The correlation coefficient does not differ significantly between subject groups.

**Figure 7:**
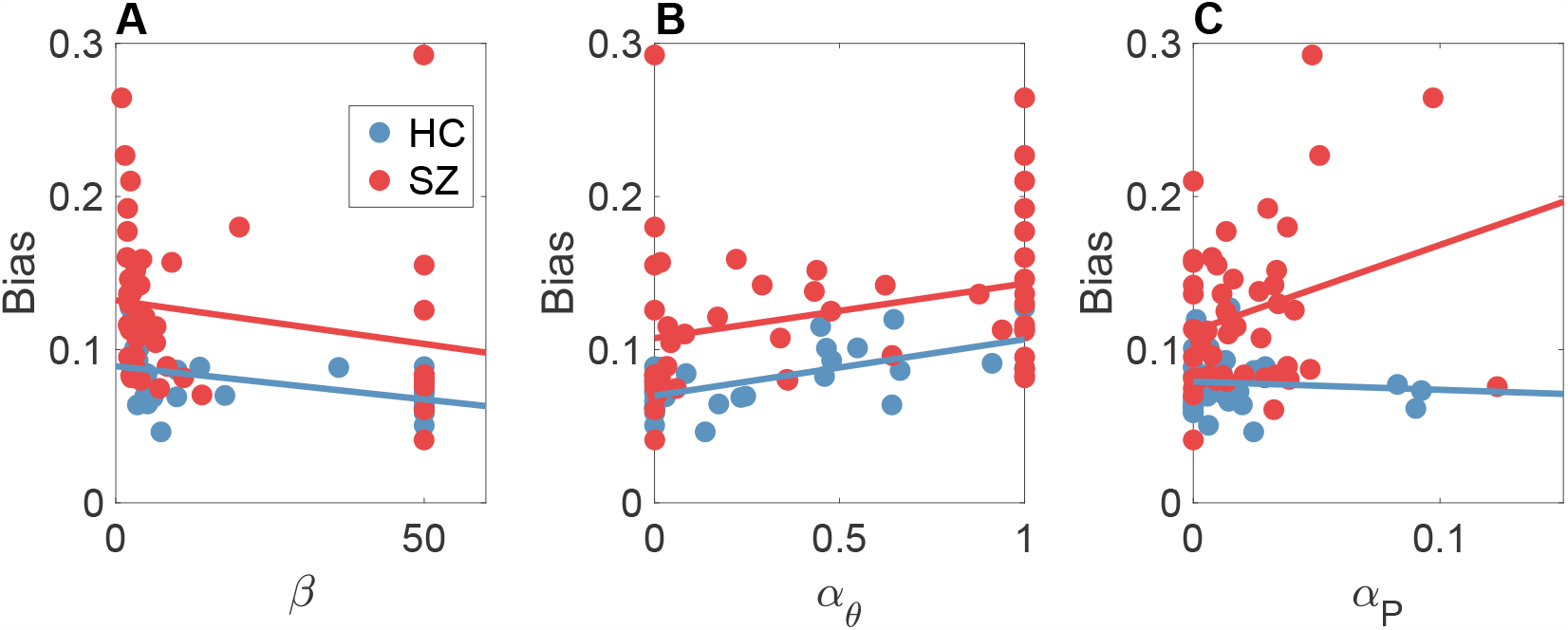
Relationship between bias and actor-critic parameters. (A) Inverse temperature. (B) Actor learning rate. (C) Marginal action probability learning rate. HC = healthy controls; SZ = schizophrenia patients.

We found a significant difference between groups for two parameters. First, the inverse temperature (*β*), which also plays the role of the capacity parameter, was higher for the healthy controls [*t*(83) = 2.29, *p <* 0.05; Figure 7A]. Second, the actor learning rate (*α*_*θ*_) was lower for the healthy controls [*t*(83) = 2.67, *p <* 0.01; Figure 7B]. There were no significant differences between groups for the critic learning rate (*α*_*V*_) or the marginal action probability learning rate (*α*_*P*_).

Putting these various observations together, we conclude that the deviation from the optimal trade-off curve exhibited by subjects (particularly those with low policy complexity) can be explained as a consequence of suboptimal learning. This suboptimality is more pronounced in the schizophrenic group, which had higher actor learning rates that in turn produced greater bias. This fits with the theoretical observation that convergence of actor-critic algorithms depends on the actor learning much more slowly than the critic (Konda and Tsitsiklis, 2000); thus, an actor that learns too fast can produce suboptimal behavior.

## Discussion

In this paper, we analyzed data from a deterministic reinforcement learning task in which the number of stimuli (the set size) varied across blocks. Both schizophrenia patients and healthy controls achieved reward-complexity trade-offs that were strongly correlated with the optimal trade-off curve, but nonetheless deviated from the optimal curve for subjects with low complexity policies. In general, schizophrenia patients had lower complexity policies and hence were more biased away from the optimal curve. However, both groups of subjects appeared to lie on the same empirical reward-complexity curve. In other words, even though the schizophrenia patients were more biased than healthy controls, they did not exhibit *excess* bias relative to the empirical curve.

One implication of this conclusion is that insensitivity to reward in schizophrenia might reflect a quasi-rational trade-off rather than a cognitive impairment *per se*. This distinction is important because it has consequences for welfare analysis and clinical interventions. If a schizophrenia patient is relatively insensitive to reward, that does not necessarily indicate that they are dysfunctional—it could alternatively reflect their preference, in which case we would not want to intervene on their decision-making processes specifically to increase reward sensitivity. On the other hand, the deviation from optimality exhibited by both healthy controls and (especially) schizophrenia patients suggests an opportunity for interventions that could improve welfare, since many individuals appear to be choosing a policy that does not maximize reward for a given resource constraint. For example, as suggested by Gershman (2020), it may be the case that individuals with lower cognitive resources may be less effective at optimization over the space of policies. Aiding this optimization process may nudge people closer to the optimal trade-off curve. How exactly does the brain solve the optimization problem? The problem is intractable for large state spaces, necessitating approximate algorithms. In particular, we formalized an actor-critic model that optimizes the cost-sensitive objective function based on trial-by-trial feedback. This model builds on earlier actor-critic models of reinforcement learning in the basal ganglia (Joel et al., 2002), and is closely related to recent cost-sensitive learning algorithms in the artificial intelligence literature (Fox et al., 2016, Grau-Moya et al., 2018, Haarnoja et al., 2018, Malloy et al., 2020). We showed that this model could account for the major features of our data. An examination of the parameter estimates revealed that the deviation from optimality could be accounted for largely by variation in the learning rate for the actor component, with larger learning rates associated with greater bias. Subjects in the schizophrenia group had both larger actor learning rates and lower inverse temperatures. This suggests that the two groups differ both in the degree of suboptimality (due to variation in the actor learning rate) and their reward-complexity trade-off (due to variation in the inverse temperature, which implicitly specifies the capacity constraint).

Unlike earlier models of memory-based reinforcement learning applied to the same data (Collins et al., 2014), the cost-sensitive actor-critic model conceptualizes memory capacity as a flexible resource rather than as a set of slots, analogous to models that have been proposed in the working memory literature (see Ma et al., 2014, for a review). Indeed, the modeling framework presented here is directly inspired by models of working memory based on rate-distortion theory and lossy compression (Bates and Jacobs, 2020, Sims et al., 2012, Sims, 2016), which formalize the trade-off between the costs and benefits of memory precision. Over the course of learning, the model adaptively compresses the policy so that it achieves the highest reward rate subject to a constraint on the average number of bits used to specify the policy. The implications of adaptive policy compression are wide-reaching: in addition to explaining quantitative aspects of choice perseveration (Gershman, 2020), it may also provide a normative explanation for different forms of action and state chunking observed experimentally (e.g., Dezfouli and Balleine, 2012, Tomov et al., 2020). Finally, the cost-sensitive actor-critic model suggests a computational rationale for the massive compression factor in the mapping from cortex to striatum (Bar-Gad et al., 2003). An important task for future work will be to assess whether these diverse phenomena can be encompassed within a single unifying framework.

As discussed in the Introduction, policy complexity is one of several forms of cognitive effort that have been studied in schizophrenia patients. Some earlier work operationalized cognitive effort in terms of task difficulty (Gold et al., 2015, Horan et al., 2015). While it is difficult to know exactly what this means in a computational sense, it is likely correlated with the duration or number of cognitive operations (what computer scientists would call computational complexity) rather than the number of bits needed to store information in memory (what computer scientists would call space complexity). The cost of optimization is another example of a time complexity cost. Thus, if the suboptimality of the empirical reward-complexity curves derives from the cost of optimization, then this would imply a relationship between computational and space complexity. We would then expect correlations between these distinct forms of cognitive effort—a hypothesis that should be pursued in future investigations.

## Acknowledgments

We are indebted to Anne Collins for making her data available. This research was supported by the Center for Brains, Minds and Machines (funded by NSF STC award CCF-1231216) and a Graduate Research Fellowship from the NSF.

## Footnotes

1 As pointed out by Culbreth et al. (2016), some of these inconsistencies may alternatively arise from the fact that earlier studies used binary choice tasks to assess cognitive demand avoidance, which may have been insufficiently sensitive to parametric variations in demand avoidance across subjects.

2 For this analysis, we made weak assumptions about the form of the empirical trade-off curve by using linear interpolation. Later, we adopt stronger parametric assumptions.

